# Compression algorithm for colored de Bruijn graphs

**DOI:** 10.1101/2023.05.12.540616

**Authors:** Amatur Rahman, Yoann Dufresne, Paul Medvedev

**Affiliations:** Department of Computer Science and Engineering, The Pennsylvania State University; Department of Biochemistry and Molecular Biology, The Pennsylvania State University; Huck Institutes of the Life Sciences, The Pennsylvania State University; Institut Pasteur, Université Paris Cité, G5 Sequence Bioinformatics, Paris, France; Institut Pasteur, Université Paris Cité, Bioinformatics and Biostatistics Hub, F-75015 Paris, France

## Abstract

A colored de Bruijn graph (also called a set of k-mer sets), is a set of k-mers with every k-mer assigned a set of colors. Colored de Bruijn graphs are used in a variety of applications, including variant calling, genome assembly, and database search. However, their size has posed a scalability challenge to algorithm developers and users. There have been numerous indexing data structures proposed that allow to store the graph compactly while supporting fast query operations. However, disk compression algorithms, which do not need to support queries on the compressed data and can thus be more space-efficient, have received little attention. The dearth of specialized compression tools has been a detriment to tool developers, tool users, and reproducibility efforts. In this paper, we develop a new tool that compresses colored de Bruijn graphs to disk, building on previous ideas for compression of k-mer sets and indexing colored de Bruijn graphs. We test our tool, called ESS-color, on various datasets, including both sequencing data and whole genomes. ESS-color achieves better compression than all evaluated tools and all datasets, with no other tool able to consistently achieve less than 44% space overhead. The software is available at http://github.com/medvedevgroup/ESSColor.

## 1 Introduction

Modern methods for analyzing biological sequences often reduce the input dataset to a set of short, fixed length strings called *k*-mers. When working with a collection of such datasets *E* = (*E*_0_, …, *E*_|*E*|−1_), it is fruitful to represent them as one union set of *k*-mers and, for each *k*-mer, the indices of the datasets to which it belongs. The set of indices of each *k*-mer is referred to as its color *class*, and *E* is referred to as a *colored de Bruijn graph* [1]. A colored de Bruijn graph (cdBG) is commonly used to represent a sequence database, such as a collection of sequencing experiments or a collection of assembled genomes. For example, it is used by tools for inferring phylogenies [2], quantification of RNA expression [3], and studying the evolution of antimicrobial resistance [4].

As sequence database sizes grow to petabytes [5], the cost of storing or transferring the data (e.g. on Amazon Web Services or in-house compute infrastructure) has underscored the need for efficient disk compression algorithms. Such costs are often prohibitive for smaller labs and, even for larger labs, limit the scale of data that can be analyzed. Large file sizes also hamper tool development, which relies on iterative loading/copying/modifying data, and reproducibility efforts, which require downloading and storing the data. For example, storing the 31-mers from 450,000 microbial genomes in compressed form takes about 12 Tb [4]. Unfortunately, there has not been a lot of work to develop methods for disk compression of colored de Bruijn graphs.

In contrast to disk compression, indexing cdBGs have received much attention [6]. A slew of data structures have been developed, optimizing various metrics such as index size, construction time, or query time (see the survey [6] and its follow up [7]). Indexing data structures exploit the structure of cdBGs and use clever tricks to compress the color classes of similar *k*-mers. But they also carry a space overhead to efficiently support queries; since this is not needed for disk compression, indexing data structures are usually not competitive with custom made cdBG compression methods.

The simplest option for compressing a cdBG is to compress each color (i.e. dataset) independently, using a compression tool designed for a single set of *k*-mers (e.g. [8]). This approach can work well when *k*-mers tend to not be shared among colors. However, most cdBGs have a large overlap between the *k*-mers of various colors. In such cases, independently compressing each color does not exploit the properties of cdBGs and, as we show in this paper, results in subpar compression ratios. There exist two tools designed specifically for disk compression of cdBGs. The first tool is unfortunately limited to only three *k*-mer sizes [9]. The second tool, called GGCAT [10] is a space efficient indexing method that, while not originally evaluated in this regard, turns out to also be a good disk compression method when combined with a generic post-compression step.

In this paper, we design, implement, and evaluate an algorithm ESS-color for the disk compression of cdBGs. We build upon the idea of spectrum-preserving string sets [11, 12, 13] and the followup compression format for a *k*-mer set [8], called *ESS*. By constructing an ESS of the union of *k*-mers in *E*, we represent the *k*-mer sequences themselves compactly. We exploit the fact that consecutive *k*-mers in an ESS have similar color classes in order to efficiently compress the color vectors of each *k*-mer.

We evaluate ESS-color on a variety of datasets, including bacteria, fungi, human, and including whole genome sequencing data, metagenome sequencing data, and whole assembled genomes. ESS-color achieves better compression than all evaluated tools and on all datasets, with all other tools using ≥ 44% more space on at least one of our datasets. On some datasets the improvement over all other tools is quite large, e.g. for a gut metagenome, all the other tools use at ≥ 27% more space than ESS-color. Compressing each color independently uses between 1.2x and 6.9x more space than ESS-color. The absolute compression ratio is more than 26x on datasets of assembled genomes and between 1.4x and 8.7x on datasets from sequencing experiments. The software is available at http://github.com/medvedevgroup/ESSColor.

## 2 Preliminaries

In this section we give some important definitions. Please refer to Figure 1 for examples of the introduced concepts.

**Figure 1:**
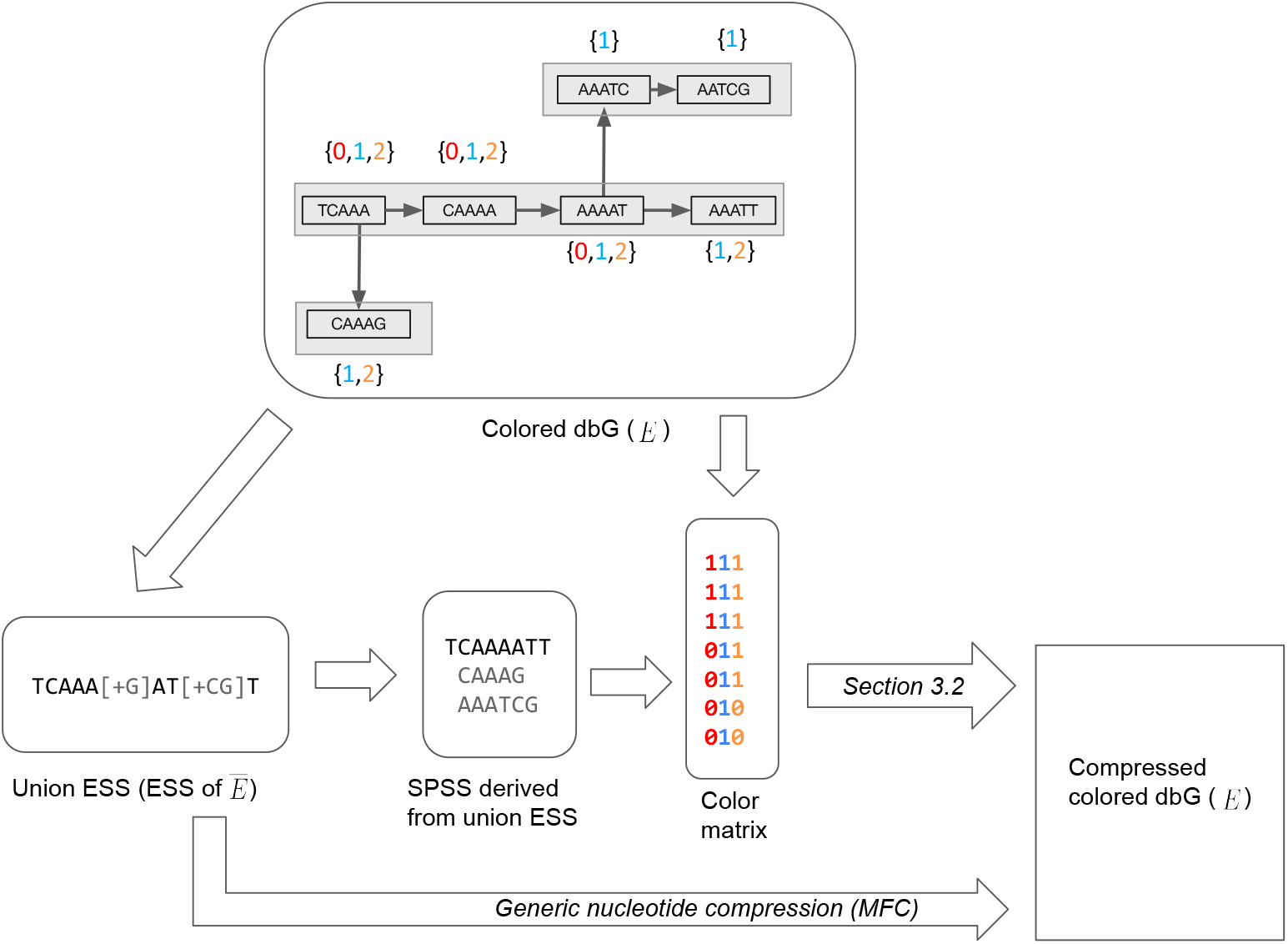
An example illustrating the various definitions in Section 2 and the first step of our compression method (Section 3.1). The input to the compression algorithm is a colored de Bruin Graph. The top panel shows an example with three colors (i.e. *C* = 3), *k* = 5, and a colored cdBG of *E* = {*E*_0_, *E*_1_, *E*_2_}. Here, *E*_0_ = {*TCAAA, CAAAA, AAAAT* }, *E*_1_ = {*TCAAA, CAAAA, AAAAT, AAATT, CAAAG, AAATC, AATCG*}, and *E*_2_ = {*TCAAA, CAAAA, AAAAT, AAATT, CAAAG*}. The color class is shown next to each *k*-mer, e.g. the color class of *TCAAA* is {0, 1, 2}. There are three distinct color classes, i.e. *M* = 3. These are {0, 1, 2}, {1, 2}, and {1}. The lower left panel shows the union ESS, i.e. the ESS representation of the set 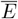 = {*TCAAA, CAAAA, AAAAT, AAATT, CAAAG, AAATC, AATCG*}. This union ESS can be decomposed into an SPSS of 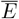, shown in the figure. The third column in bottom panel shows the color matrix, with *k*-mers in the order of the SPSS. To obtain the final compressed representation the color matrix is compressed using the algorithm we describe in Section 3.2.

### Strings

A string of length *k* is called a *k-mer*. We assume *k*-mers are over the DNA alphabet. A string *x* is canonical if it is the lexicographically smaller of *x* and its reverse complement. Let *K* be a set of *k*-mers. A *spectrum-preserving string set (SPSS)* of *K* is a set of strings *S* such that each string *s* ∈ *S* is at least *k* characters long, every *k*-mer that appears in *S* appears exactly once, and the set of *k*-mers that appear in *S* is *K* [11, 12, 13]. For example, if *K* = {*ACG, CGT, CGA*}, then {*ACGT, CGA*} would an SPSS of *K*. Note that *K* can have multiple spectrum-preserving string sets. There are several efficient tools for computing an SPSS so as to minimize the total number of characters [14, 10]. In this paper, we rely on the implementation in [8]. Each string in the resulting SPSS is referred to as a *simplitig*.

### Compression of a *k*-mer set

ESS is a disk-compression format to store a set of *k*-mers *K*. It was introduced in [8] as the output of a compression tool, which, in this paper, we will refer to as ESS-basic. The technical details of the format and of the tool are irrelevant for this paper and can be viewed as black boxes. An ESS representation cannot be queried efficiently but can be decompressed into an SPSS of *K*. This output gives an ordering of the *k*-mers of *K*, and therefore the ESS compression of *K* induces an ordering on *K*. Note that because the decompression algorithm is deterministic, by storing an ESS representation, we are implicitly storing an SPSS representation as well.

### Colored *k*-mer sets

Let *C* > 0 be an integer indicating the number of colors. Let *E* = {*E*_0_, …, *E*_*C*−1_} be a set of *C k*-mer sets, also referred to as a *colored de Bruijn graph*. Let 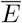 be the set of all *k*-mers in *E*, i.e. 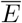 = {*x* | ∃*i* s. t. *x* ∈ *E*_*i*_}. The *(color) class* of a *k*-mer *x* ∈ 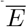 is the set of indices *i* such that *x* ∈ *E*_*i*_. The *color vector* of *x* is a binary vector of length *C* where position *i* is 1 iff *x* ∈ *E*_*i*_.

### Non-compressed representation of cdBGs

Assume you have an ordering of 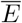, e.g. the one given by an ESS of 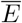. A *color matrix* of *E* is a file with row *i* containing the color vector of the *i*^th^ *k*-mer. Storing an ESS of 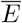 together with a color matrix of *E* is a lossless representation of *E*.

## 3 Methods

In this section, we describe our algorithm ESS-color for the compression of cdBGs. Let *E* = {*E*_0_, …, *E*_*C*−1_} be a colored dBG over *C* colors. Recall that 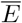 is the set of all *k*-mers in *E*. Let *M* denote the number of distinct color classes in *E*.

ESS-color can accept input in one of two formats. First, it can accept each *E*_*i*_ stored in the KFF file format [15]. Alternatively, we can take as input a collection of FASTA files, each one assigned one of *C* colors, and an abundance parameter *a. E*_*i*_ is then implicitly defined as the set of all canonical *k*-mers that appear at least *a* times in the FASTA files of color *i*. We obtain *E*_*i*_ by running KMC [16] on the FASTA files of color *i*.

### 3.1 Building a color matrix of *E* and compressing 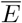

In this step, we first compress 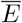 using ESS-basic [8] and then build the color matrix of *E*, ordered by the ESS order. Specifically, we first compress the nucleotide sequences themselves, i.e. we run ESS-basic [8] on all the input files jointly. We refer to this as the *union ESS*. We then decompress this file to obtain an SPSS of 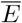, denoted by *S*. From *S*, we build an SSHash dictionary [17] that allows us to map each *k*-mer in 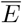 to its rank in *S*. We then build on top of the KMC API to read in the binary files representing *E*_0_, …, *E*_*C*−1_ and output a color matrix ordered by the SSHash dictionary. At the end of this stage, we have the union ESS, which is retained in the final compression output, and we have *S* and the color matrix, which are used in later stages but not retained in the final compression output.

### 3.2 Compression of the color matrix

Given an SPSS *S* of 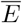 and a color matrix of *E* over the order induced by *S*, we now generate a compressed representation of the matrix. Our representation consists of a *global class table* and, for every simplitig of *S*, a few bits of metadata, a *local class table* and one bitvector *m*. The local class table is optional, as we describe below. Figure 2 gives a schematic representation. We now explain each of these in detail.

**Figure 2:**
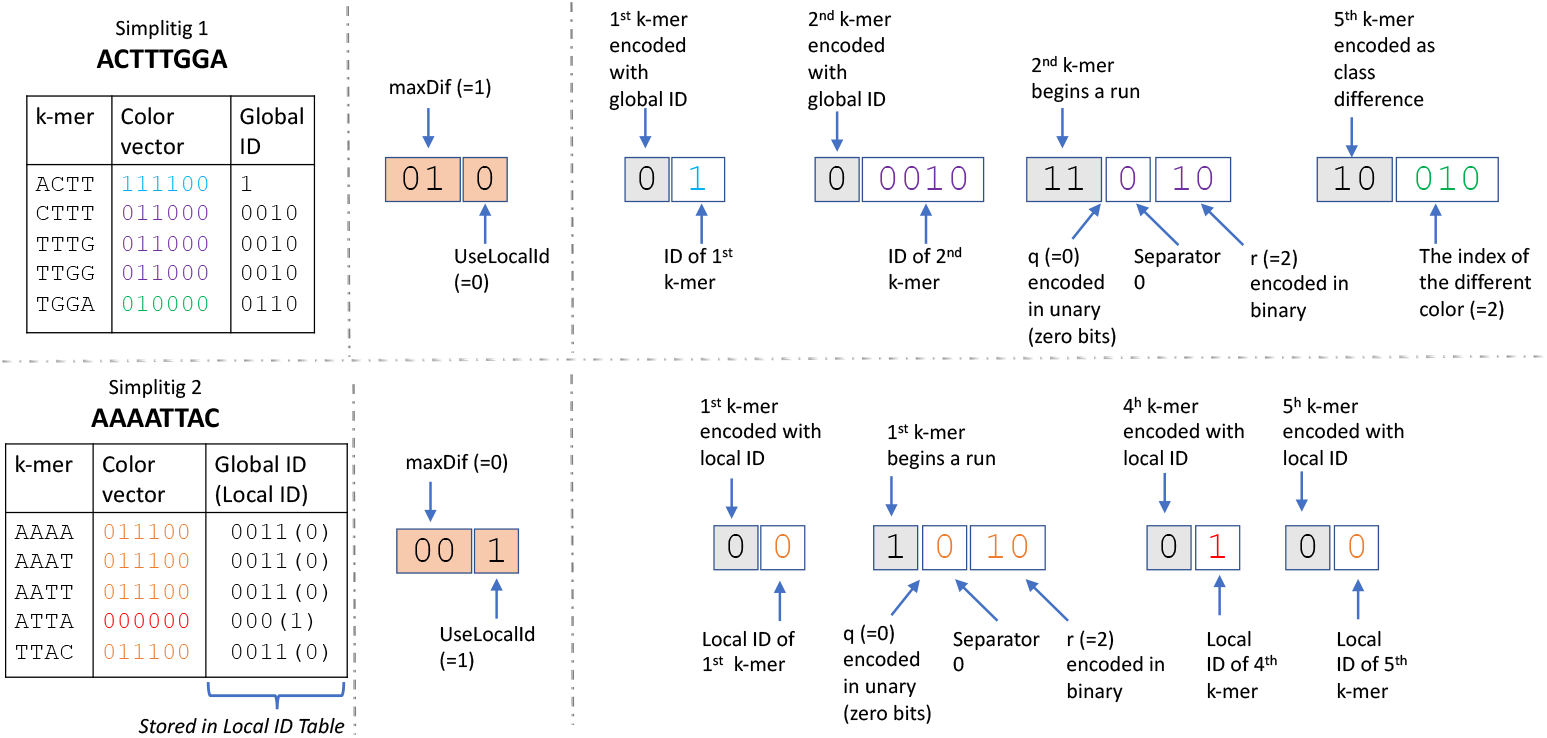
Example of how we compress the color matrix (Section 3.2). The top panel shows the compression of a simplitig ACTTTGGA and the bottom panel shows the compression of a simplitig AAAATTAC. Other simplitigs exist but are not shown. In this example, *C* = 6, *k* = 4, and *runDivisor* = 4. The figure is divided into three columns. The first column shows the information that the algorithm has about each *k*-mer (i.e. its color vector and corresponding global ID). The second column shows the metadata which holds the values of *maxDif* and *useLocalId* for the corresponding simplitig. The third column shows the *m* vector. Within the *m* vector, the type of encoding is shown in gray colored boxes and other values are shown in white boxes. The colors of the values inside the white boxes correspond to the color used for the corresponding *k*-mers’ color vector in the first column of the figure. Note that in the bottom simplitig, a local table is used. In particular, there are two distinct color vectors in this simplitig, with global ID 0011 assigned to local ID 0 and global ID 000 assigned to local ID 1.

#### Global class table

> For most applications, the number of distinct color vectors *M* is significantly smaller than 2^*C*^. Hence, the color matrix representation, which uses *C* bits per *k*-mer, is very inefficient. Instead, we use Huffman coding to assign a *global ID* to each class, so as to minimize the number of bits that will be used to store these IDs later (this is similar to what was done in [18]). To do this, we scan the color matrix to determine all the distinct classes and the number of *k*-mers that have each class. We then use Huffman coding to assign a global ID to each distinct class, so that more frequent global IDs tend to use less bits. This table is then stored in two forms: one that is compressed to disk, and the other that is stored in memory to be used during the compression algorithm.
>
> We store the table on disk using three files: a color encoding Δ, a boundary bitvector *b*, and a text file. First, we sort the color classes in increasing numerical order, interpreting each color vector as a *C*-bit integer. For Δ, we write a concatenation of the *M* color vectors to disk, with the first color vector being written using *C* bits and the following colors being encoded as a difference with their predecessor. Specifically, if *h*_*i*_ is the Hamming distance between the *i*^th^ and the (*i* − 1)^st^ color vectors, then we use *h*_*i*_⌈log *C*⌉ bits to encode the indices where the *i*^th^ color vector is different from the (*i* − 1)^st^ color vector. We also store a boundary bitvector *b* which is the same length as Δ and contains a ‘1’ whenever Δ starts a new color class. Finally, we store the frequencies of the color classes in a text file. These three files are then sufficient to reconstruct the global IDs during decompression. ^1^
>
> Simultaneously, we need to be able to map a color vector to an ID during the compression process. To do this, we create a minimal perfect hash function *h* (CHM [20]) that maps from each distinct color vector to an integer between 0 and *M* − 1. We then maintain an array *A* of size *M*, where for each color vector *c, A*(*h*(*c*)) holds the global ID of color vector *c*.

After the global class table is created, we process the simplitigs of *S* one at time. For each simplitig, we dynamically set two parameters: a boolean variable *UseLocalID*, and an integer 0 ≤ *maxDif* ≤ 2. We postpone the discussion on how these are set until the end of the section. The values of *maxDif* and *UseLocalID* are stored using 3 bits of metadata per simplitig. If *UseLocalID* is set, we create a local class table:

#### Local class table

> In the case that the frequencies of color classes are evenly distributed, we need approximately log *M* bits to represent the global class ID of each *k*-mer. We observe that sometimes a class is used at multiple locations of a simplitig, in which case using log *M* bits for each occurrence can be wasteful. Let *ℓ* be the number of distinct classes appearing in a simplitig. To save space on class IDs, we create a separate *local class table*, which maps from *ℓ* integers, called *local class IDs*, to their respective global IDs. Then, the encoding of *k*-mer classes for this simplitig can use local class IDs, which take only log *ℓ* space. The local class table is written to disk, with log *M* bits encoding *ℓ* followed by *ℓ* consecutive global class IDs together

The bitvector *m* is constructed by scanning the simplitig from left to right and, for each *k*-mer *x*, deciding how to encode it, and appending that encoding to *m*. Intuitively, the encodings follow three basic possibilities. The first possibility is to just append *m* with the *k*-mer’s class ID. Second, we observed in practice that simplitigs often contain runs of *k*-mers with identical classes, in which case we can append *m* with the length of the run, rather than writing out each class IDs (such runs are similar to the monotigs of [21]). Finally, we often observe that a *k*-mer has a color vector with a small Hamming distance (i.e. 1 or 2) to that of the previous *k*-mer. In this case, we append *m* with the indices in the color vector that are different. Since there are three types of encoding, we will also need to prepend each encoding with two bits denoting the type of encoding. Formally, for each *k*-mer in a simplitig, we choose one of four options:

#### Skip

> This option is invoked if *x* is not the first or last *k*-mer in its simplitig and has the same class as the preceding and succeeding *k*-mer. In this option, nothing is appended to *m*.

#### Small class difference

> Let *h* be the Hamming distance between the color vector of *x* and the color vector of the preceding *k*-mer in the simplitig. This option is invoked when 0 *< h* ≤ *maxDif*. First, we append *m* with ‘10’ to indicate that the following encoding will encode a class difference. If *maxDif* = 2, then we append *m* with a ‘1’ to indicate that *h* = 2 or a ‘0’ to indicate that *h* = 1. If *maxDif* = 1, then we do not append this extra bit, since it is implicit. Then, we append *m* with *h* log *C* bits which list the colors that are different. Note that setting *maxDif* = 0 effectively disables this type of encoding.

#### End of run

> This option is invoked if *x* has the same class as the preceding *k*-mer and either has a different class than the succeeding *k*-mer or is the last *k*-mer in the simplitig. First, we indicate that the following encoding will encode a run length by appending *m* with ‘11’ if *maxDif* > 0 and ‘1’ if *maxDif* = 0. This difference is due to the fact that if *maxDif* = 0, then there are only two types of encodings and so we can just use one bit for the type.
>
> Let *runLen* be the number of consecutive *k*-mers that preceded *x* (not including *x*) and have the same class. We encode *runLen* by separating it into a quotient *q* and remainder *r* (with respect to a global parameter *runDivisor*), and then encoding the quotient *q* in unary and the remainder *r* in binary. Formally, let 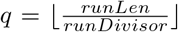 and *r* = *runLen* mod *runDivisor*. We append to *m q* ‘1’s followed by a ‘0’. Then, we append to *m* the binary encoding of *r*, using log *runDivisor* bits. For example, if *runDivisor* = 16 and *runLen* = 21, then *q* = 1 and *r* = 5, and *m* is appended with 100101. Observe that a smaller value of *runDivisor* results in more bits used to encode long runs (i.e. *q* is larger) while a larger value of *runDivisor* uses more bits to encode short runs (i.e. log *runDivisor* is larger). We found that a default value of 16 works best in our experiments.

#### Store the class ID

> This option is invoked when none of the criteria for the other options are satisfied. In this case, we append *m* with ‘0’ to indicate the type of encoding, followed by the class ID of *x*. If *UseLocalID* is set, we use the local class ID, otherwise we use the global class ID.

After finishing with all simplitigs, we compress the global class table, local class tables, and *m* using RRR [22] and write them to disk.

Setting the parameters *UseLocalID* and *maxDif* involves trade-offs that are difficult to quantify in advance. For example, the cost of having to store the local class table may exceed the benefits of using less bits to encode class IDs for a simplitig where every present class ID is contained within a single run. Similarly, when *d* is too large, then writing the positions of the color differences to *m* can take more space then just writing the class ID. Moreover, there is a benefit of setting *d* = 0, since it enables to save one bit per run by using ‘1’ instead of ‘11’ for the ‘end of run’ encoding. All bitvectors are additionally compressed with RRR, making it difficult to determine in advance which parameters result in the least space. We therefore try all possibilities of *maxDif* ∈ {0, 1, 2} and *UseLocalID* ∈ {*True, False*}, and, for each combination, compute the encoding. We then use the encoding that takes less space and disregard the rest. Though this step can likely be optimized, we found that the time taken to try all possibilities was not a large factor in the overall compression time.

The decompression algorithm for the *m* vector is straightforward since our color matrix compression scheme is designed to be unambiguously decompressed. Simultaneously, we decompress *E* with ESS-decompress. The result is an SPSS *S* of 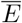 and a color matrix of *E* in the order of *S*. If the output is to be processed downstream in a streaming manner, our decompression algorithm can trivially stream out *k*-mer sequence and color vectors, one *k*-mer at a time.

## 4 Results

### 4.1 Evaluated tools and datasets

As far as we are aware, there are two other tools designed for compressing colored de Bruijn graphs: KS [9] and GGCAT [10]. We refer to the first tool as KS after the authors’ last names [9]. KS is limited to support only three *k* values (15, 19, and 23), so we compare against it for *k* = 23 but also evaluate ESS-color on a more practical *k* value of *k* = 31. For GGCAT, we additionally compressed its Fasta output file with MFC and its binary color table with gzip to maximize its compression ratio. We also compare ESS-color against the naive approach of compressing each color independently using the algorithm of [8], which we refer to as *ESS-basic*.

Table 1 shows the datasets we use for evaluation and their properties. We chose five datasets so as to cover a broad range of input types. Three of the datasets are from assembled genomes, one is from RNA-seq reads, and one is from metagenome reads. We used all *k*-mers from the three assemblies datasets and all *k*-mers that appear at least twice from the two read datasets. The datasets cover various species, from Bacteria to Fungi to Human. Concretely, we have 1) 100 arbitrarily selected *E*.*coli* strains from GenBank, 2) an arbitrary subset of 10 of those, 3) 20 arbitrarily selected fungi sequences from RefSeq, 4) gut microbiome read sets from nine individuals sequenced in [23], and 5) 19 paired-end, human, bulk RNA-seq short-read experiments previously used in [19]. All accession numbers are listed in https://github.com/medvedevgroup/ESSColor/wiki/Experiments.

**Table 1:** Dataset characteristics. *C* is the number of colors and *M* is the number color classes.

| Source | type | C | M | n. $k$-mers $\times 10^6$<br>(% single color) | | n. simplitigs $\times 10^6$ | |
| --- | --- | --- | --- | --- | --- | --- | --- |
| | | | | $k = 23$ | $k = 31$ | $k = 23$ | $k = 31$ |
| E. coli | assemblies | 100 | 542,545 | 27 (30%) | 31 (31%) | 0.5 | 0.5 |
| E. coli | assemblies | 10 | 826 | 13 (38%) | 14 (41%) | 0.2 | 0.2 |
| Fungi | assemblies | 20 | 13,227 | 394 (93%) | 409 (93%) | 1.8 | 1.7 |
| Gut | metagenome reads | 9 | 511 | 2,236 (67%) | 2,477 (70%) | 76 | 95 |
| Human | RNA-seq reads | 19 | 9,654 | 120 (71%) | 103 (75%) | 7.2 | 10 |

Finally, all experiments were run on a server with an Intel(R) Xeon(R) CPU E5-2683 v4 @ 2.10GHz processor with 64 cores and 512 GB of memory. We ran all tools in unrestricted memory mode. We used 8 threads for all tools and their components, whenever they supported multi-threading.

### 4.2 Comparison against other disk compression tools

Table 2 shows the bits per *k*-mer achieved by ESS-color compared with KS, GGCAT ESS-basic, and a p7zip compression of the original fasta file. ESS-color achieves better compression than all evaluated tools and on all datasets. No other tool was able to consistently achieve less than 44% space overhead compared to ESS-color. On some datasets the improvement over all other tools is quite large, e.g. for Gut (*k* = 23), all the other tools use at least 27% more space than ESS-color.

**Table 2:** Compression results, in bits per *k*-mer. ESS-color is our new tool. ESS-basic is the non-integrative approach of compressing every color separately. KS is tool from [9], and a hyphen indicates that it does not support *k* > 23. fa.p7zip is the p7zip compression of the original data. We show bits per *k*-mer, which is the total compressed size divided by the number of distinct *k*-mer in the input (i.e. |*E*|). Compression ratios can be inferred by comparing to the fa.p7zip column.

| Dataset | $k = 23$ | | | | | $k = 31$ | | | | |
| --- | --- | --- | --- | --- | --- | --- | --- | --- | --- | --- |
|  | fa.p7zip | ESS-color | KS | GGCAT | ESS-basic | fa.p7zip | ESS-color | KS | GGCAT | ESS-basic |
| Ecoli100 | 37 | 5.2 | 17.3 | 6.4 | 35.0 | 33 | 4.4 | – | 5.5 | 30.3 |
| Ecoli10 | 8 | 2.9 | 5.3 | 3.3 | 7.3 | 7 | 2.6 | – | 3.0 | 6.6 |
| Fungi20 | 2.4 | 2.2 | 2.5 | 2.3 | 2.3 | 2.3 | 2.2 | – | 2.3 | 2.5 |
| Gut9 | 60 | 3.7 | 5.7 | 4.9 | 4.7 | 54 | 3.4 | – | 4.9 | 4.4 |
| HumRNA19 | 112 | 5.6 | 8.8 | 7.1 | 10.4 | 131 | 8.9 | – | 10.5 | 12.1 |

The compression ratio of ESS-color relative to the original Fa.p7zip files varies (Table 2). For the read datasets, it is more than 15x, since high-coverage FASTA files are by their nature very redundant. For the assemblies datasets, the ratio is between 1x and 7x. We found that a good predictor of compressibility of assemblies is the percentage of *k*-mers that have exactly one color (Table 1). At one extreme, 93% of Fungi20 *k*-mers have exactly one color, and ESS-color achieves little improvement over fa.p7zip. At the other extreme, only 30% of Ecoli100 *k*-mers have exactly one color, and the compression ratio is relatively high at 7.1x (for *k* = 23). This trend makes intuitive sense since single-color *k*-mers do not benefit from ESS-color’s multi-color compression algorithm.

KS is not as effective as ESS-color on our datasets, using between 1.4x and 3.3x more space than ESS-color (Table 2). We note that even though KS is also designed to exploit the fact that *k*-mers are shared across colors, it makes a different design trade-off compared to ESS-color. Specifically, it does not allow simplitigs to extend beyond a single color class (resulting in more space needed to store *k*-mers), but, in exchange, it is more efficient in storing color information.

GGCAT is generally the closest competitor against ESS-color, using between 14 and 44% more space on the non-Fungi datasets (note that for Fungi all tools did well). Like ESS-color, it builds a kind of global class table, constructs an SPSS of the *k*-mers, and annotates each run of single-class *k*-mers with their class ID. Unlike ESS-color, however, it does not use ESS, does not encode small class differences, and does not use local class tables.

As expected, ESS-basic is not as effective as ESS-color, using up to 6.9x more space than ESS-color (Table 2). These results are not surprising since ESS-basic does not exploit the redundancy created by shared *k*-mers across samples. For the assemblies datasets, the compression improvement of ESS-color over ESS-basic closely tracks that of ESS-color over the original fa.p7zip. For the sequencing datasets, ESS-basic uses between 1.3 and 1.9 more space than ESS-color.

Tables 3 and 4 show the run time and memory usage of compression, respectively. Here, ESS-color is outperformed by other tools. In particular, if optimal compression space is not needed, then GGCAT becomes a good alternative to ESS-color. Note that the decompression time (Table 5) is an order of magnitude smaller compared to the compression times.

**Table 3:**
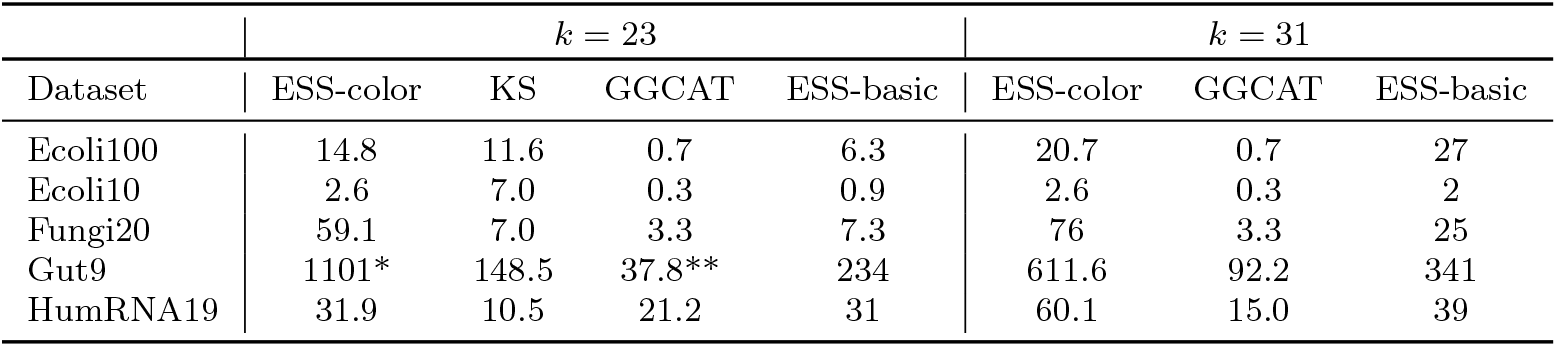
Time (min) used by the various compression algorithms. For the Gut9 ESS-color run with *k* = 23 (marked with *), we used an unoptimized implementation of the color matrix generation step, since SSHash was not working as expected. For GGCAT on Gut9 with *k* = 23 (denoted by **), the original run crashed because of exceeding the number of open files allowed by the operating system. We therefore re-ran GGCAT using our simplitigs as a starting point, which allowed the run to complete. However, the time shown here does not include the time we used to construct the simplitigs.

**Table 4:** Maximum memory (Gb) used by the various compression algorithms. The (*) and (**) annotations are the same as in Table 3.

| | $k = 23$ | | | | $k = 31$ | | |
| --- | --- | --- | --- | --- | --- | --- | --- |
| Dataset | ESS-color | KS | GGCAT | ESS-basic | ESS-color | GGCAT | ESS-basic |
| Ecoli100 | 1.2 | 0.9 | 1.5 | 1.2 | 1.1 | 1.4 | 1 |
| Ecoli10 | 0.6 | 3.2 | 0.8 | 1.1 | 0.6 | 0.8 | 1 |
| Fungi20 | 5.4 | 3.2 | 4.8 | 5.9 | 4.3 | 3.9 | 6 |
| Gut9 | 174.6* | 50.6 | 87.2** | 33.2 | 121 | 78.2 | 119 |
| HumRNA19 | 26.8 | 6.0 | 8.8 | 9.1 | 12.1 | 10.2 | 7 |

**Table 5:** Time and memory for decompression of ESS-color, for *k* = 31.

| Dataset | Memory (GB) | Time (min) |
| --- | --- | --- |
| Ecoli100 | 0.5 | 2 |
| Ecoli10 | 0.5 | 1 |
| Fungi20 | 0.5 | 13 |
| Gut9 | 8.5 | 90 |
| HumRNA19 | 1.0 | 5 |

### 4.3 Inside the space usage of ESS-color

ESS-color’s compressed representation includes several components, with the major ones being the union ESS, the *m* vector, the global table, and the local tables. Table 6 shows that the majority of space used by ESS-color is taken by the union ESS. Except for Ecoli100, the rest of the space is taken up almost exclusively by *m*. For Ecoli100, which has the largest number of colors, the global table takes 23% of the total space.

**Table 6:**
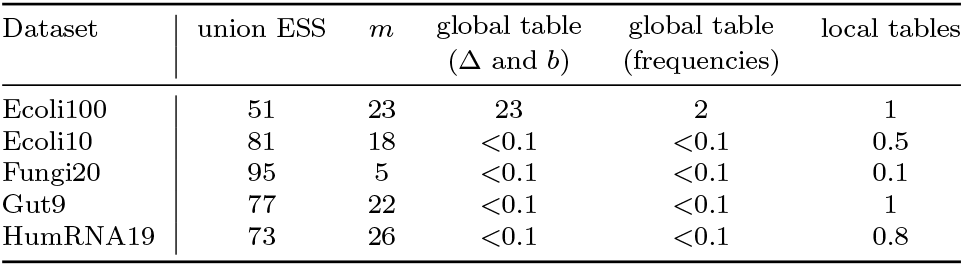
Breakdown of the space usage (in percentage of total space) of the components of ESS-color, for *k* = 31. Note that all components except union ESS are shown after compression with RRR.

| Dataset | union ESS | $m$ | global table<br>($\Delta$ and $b$) | global table<br>(frequencies) | local tables |
| --- | --- | --- | --- | --- | --- |
| Ecoli100 | 51 | 23 | 23 | 2 | 1 |
| Ecoli10 | 81 | 18 | <0.1 | <0.1 | 0.5 |
| Fungi20 | 95 | 5 | <0.1 | <0.1 | 0.1 |
| Gut9 | 77 | 22 | <0.1 | <0.1 | 1 |
| HumRNA19 | 73 | 26 | <0.1 | <0.1 | 0.8 |

Recall that ESS-color chooses one of six different compression modes for each simplitig, i.e. *UseLocalId* ∈ {0, 1} and *maxDif* ∈ {0, 1, 2}. In order to access the relative contribution of the various compression techniques, we count the frequency with which each mode occurs (Table 7). First, we observe that the idea of a local table was rarely helpful on our data. Local tables are only beneficial when a single color class appears in more than one run in a simplitig, which apparently was rare. Second, the majority of simplitigs use *maxDif* = 0. This mode is optimal when the simplitig has just one color class. There is also a more complicated trade-off since setting *maxDif* > 0 adds one extra bit for each run encoding, which may outweigh the benefits of encoding some *k*-mers with a class difference. Third, the Gut dataset demonstrates the benefit of encoding class differences, especially compared to GGCAT. It is the dataset with the highest percentage of simplitigs using *maxDif* > 0 (21%) and, simultaneously, it is also the dataset where GGCAT uses the most space relative to ESS-color (44%).

**Table 7:** The percentage of simplitigs (*k* = 31) that fall into the six compression modes, i.e. combinations of *UseLocalID* and *maxDif*.

| Dataset | $UseLocalID = \text{False}$ | | | $UseLocalID = \text{True}$ | | |
| --- | --- | --- | --- | --- | --- | --- |
| | $maxDif = 0$ | $maxDif = 1$ | $maxDif = 2$ | $maxDif = 0$ | $maxDif = 1$ | $maxDif = 2$ |
| Ecoli100 | 75 | 20 | 5 | <0.1 | <0.1 | <0.1 |
| Ecoli10 | 82 | 16 | 2 | <0.1 | <0.1 | <0.1 |
| Fungi20 | 99 | 1 | 0.2 | <0.1 | <0.1 | <0.1 |
| Gut9 | 80 | 19 | 2 | 0.2 | <0.1 | <0.1 |
| HumRNA19 | 91 | 9 | 0.1 | 0.1 | <0.1 | <0.1 |

### 4.4 Comparison to indexing data structures

There exist numerous indexing data structures for cdBGs [6]. Indexing data structures are similar to disk compression but additionally support efficient membership and color queries. We expect this overhead to make them non-competitive with respect to disk compression schemes. To verify this, we compared the space taken by ESS-color against three indexing approaches. We note that since these approaches are designed for indexing, they do not implement decompression and are thus not viable for disk compression in their current state. We also note that GGCAT also supports indexing, but, since it is trivial to decompress, we included it in the main analysis of Section 4.2.

The first two approaches are ones that are shown in [24, 25] to be the most space efficient. These are RowDiff+, which is the latest version [25] of RowDiff [24], and Rainbow-MST [24], which is a space-improved version of Mantis-MST [19]. As a trivial improvement, we further compress these indices using gzip. The third approach we compare to is a natural hybrid of ESS-color and the RowDiff indexing algorithm for cdBGs [24]. We refer to this as *RowDiff-ESS* and describe it in detail in the Appendix. We do not compare against other indices such as REINDEER [21], Bifrost [26], Themisto [27], or Mantis-MST [19], because they are less space efficient than RowDiff+, and we do not compare against Sequence Bloom Tree approaches (e.g. [28, 29]) because they are lossy.

Table 8 shows the results. As expected, the compression ratios of these indexing tools are not competitive against ESS-color. Even the most space efficient indexing approach for each dataset takes 60% more space than ESS-color. We do note that GGCAT, which was shown in Table 2, is an exception, since it implements both efficient indexing and disk compression.

**Table 8:** Compression results in bits per *k*-mer (*k* = 31) of indexing approaches, compared to ESS-color.

| Dataset | ESS-color | RowDiff-ESS | RowDiff+ | Rainbow-MST |
| --- | --- | --- | --- | --- |
| Ecoli100 | 4.4 | 14.6 | 8.0 | 34.2 |
| Ecoli10 | 2.6 | 7.8 | 6.3 | 19.3 |
| Fungi20 | 2.2 | 3.6 | 4.3 | 9.5 |
| Gut9 | 3.4 | 11.2 | 40.6 | 15.6 |
| HumRNA19 | 8.9 | 37.9 | 112.6 | 19.1 |

## 5 Conclusion

Colored de Bruijn graphs are a popular way to represent sequence databases. In spite of their ever-growing sizes, there have not been many specialized tools for compressing them to disk. In this paper, we present a novel disk compression algorithm tailormade for cdBGs that achieves superior space compression compared to all other tools on the evaluated datasets.

Our algorithm is a novel combination of ideas borrowed from previous work on disk compression of *k*-mer sets and indexing of cdBGs. We use a spectrum-preserving string set (SPSS) as a basis for both compressing the nucleotide sequences and for ordering the rows in the color matrix. By using the SPSS ordering, we can avoid the costly storage of an indexed de Bruijn graph (e.g. BOSS in [24, 25] or a counting quotient filter in [19]). We also exploit the fact that consecutive *k*-mers in an SPSS likely have the same or similar color class. A major component of our approach is that we select a different compression scheme for each simplitig, depending on what gives the best compression on that simplitig.

The most important practical direction for future work is to improve the running time of our algorithm. The generation of the union ESS is done by ESS-basic. ESS-basic can be sped up by extending the latest SPSS generation tools [14, 10] to also compute an ESS. We could even build on top of GGCAT, taking advantage of their efficient implementation (unfortunately, GGCAT was only released once our project was near completion). Another bottleneck is the color matrix generation step, which could be parallelized or even avoided by using color lists.

A theoretical future direction is to derive bounds on the bits used by the compression scheme. Unfortunately, we do not see an easy way to do this, since the choice of encoding depends on the order of the *k*-mers in the SPSS and on the decomposition of the *k*-mers into simplitigs. It is unclear to us how to capture these properties as a function of the input data.

Finally, we could further improve the compression algorithm by modifying the SPSS generated by the ESS-basic algorithm. Currently, the choice of how to select from multiple simplitig extensions is made arbitrarily. Instead, the choice could be made to use the extension that has the most similar color class. Such a modification to the SPSS construction algorithm would likely be computationally non-trivial, since it would require accessing the color information.

## Acknowledgements

We would like to thank R. Chikhi for helpful discussions. This material is based upon work supported by the National Science Foundation under grant nos. 2138585 and 1931531. Research reported in this publication was also supported by the National Institutes of Health under Grant NIH R01GM146462 (to P.M.). A.R. was supported by the National Institutes of Health Computation, Bioinformatics, and Statistics (CBIOS) training program. The content is solely the responsibility of the authors and does not necessarily represent the official views of the National Institutes of Health.

## 6 Appendix

Here we describe *RowDiff-ESS*, the hybrid of ESS-color and the RowDiff indexing algorithm for cdBGs [24]. Though the approach turned out to not be comptetive against ESS-color, we describe it here for completeness. The RowDiff index is composed of two parts: BOSS, which is an index of 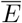, and a compressed color matrix whose labels are implicitly given by BOSS. Because of its structural similarity to our approach, we can swap out BOSS (which supports queries and is therefore not space efficient for disk compression) with an ESS of 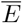. We then feed the *k*-mer ordering implied by the ESS to the RowDiff color matrix compression algorithm. The space used by this scheme is the ESS space plus the space of the RowDiff’s color matrix, compressed with gzip.

## Footnotes

1 For readers familiar with Mantis-MST [19], we also tried their approach for storing our global table. Surprisingly, we found that our approach outperformed their more sophisticated approach, at least in our datasets. Though the Mantis-MST approach resulted in a smaller Δ vector, the overhead of storing the tree parent vector outweighed this gain.

